# Kupffer cells dictate hepatic responses to the atherogenic dyslipidemic insult

**DOI:** 10.1101/2022.10.13.512086

**Authors:** Sanna Hellberg, Osman Ahmed, Yuyang Zhang, Giada Di Nunzio, Hanna M Björck, Roy Francis, Linn Fagerberg, Anton Gisterå, Xueming Zhang, Jari Metso, Valentina Manfé, Yosdel Soto, Anders Franco-Cereceda, Per Eriksson, Matti Jauhiainen, Peder S. Olofsson, Stephen G. Malin

**Author notes:** These authors contributed equally.

## Abstract

Apolipoprotein-B (APOB) containing lipoproteins are causative for atherosclerotic cardiovascular disease. Whether the vasculature is the initial responding site or if atherogenic-dyslipidemia effects other organs simultaneously is unknown. We set out to discover how the liver responds to a dyslipidemic insult through the creation of inducible mouse models based on human familial hypercholesterolemia mutations and *in vivo* tracing of APOB. An acute transition to atherogenic APOB-lipoprotein plasma levels resulted in rapid accumulation of triglycerides and cholesterol in the liver. Single cell RNA-seq and flow cytometry disclosed that multiple immune cells have the ability to engulf APOB-lipoproteins. However bulk RNA-seq of the liver revealed an inflammatory Kupffer cell-specific transcriptional program that could not be activated by a western diet alone. Depletion of Kupffer cells through clodronate liposomes or CD8 T cell targeting rapidly raised plasma lipoprotein levels, indicating that these liver macrophages help restrain and buffer atherogenic lipoproteins, whilst simultaneously secreting pro-atherosclerotic factors into plasma. Our results place Kupffer cells as a key gateway in organizing systemic responses at the initiation of atherosclerosis.

## Introduction

APOB-lipoproteins, most notably low-density lipoprotein (LDL), are the causal agents of atherosclerosis, due to their ability to be retained, modified and engulfed at susceptible sites in the vasculature(1, 2). The liver is central to atherosclerotic cardiovascular disease (ACVD) due to its primary role in APOB-lipoprotein production and clearance by hepatocytes (3). Non-parenchymal Liver cells, especially the resident macrophage Kupffer cells (KCs), take up significant amounts of APOB-lipoproteins in rats (4), rabbits (5) humans(6) and in the specific context of hypercholesterolemic mice with hereditary haemochromatosis (7). However, the function of the liver and its specific cells at the initiation of atherosclerosis remains unexplored.

In addition to their well-recognized role in initiating and sustaining atherosclerosis, APOB-lipoproteins can also contribute to pathologies located in other organs, including the liver itself (8, 9). Non-alcoholic fatty liver disease (NAFLD) describes a range of liver disorders that is thought to start with simple lipid accumulation – steatosis – through to irreversible cirrhosis(10). Free fatty acids released by dysfunctional adipose tissue have traditionally been the focus of how lipids contribute to NAFLD. This simplified view has been challenged by the contributions of APOB-lipoproteins to the disease process. NAFLD and atherosclerosis also share comorbidities including diabetes, and both can be associated with elevated levels of triglycerides and remnant APOB-lipoproteins. However, NAFLD confers an increased risk of ACVD beyond the sum of these individual components in a manner that is poorly understood (11, 12). Atherosclerosis and NAFLD may also share similar mechanisms of disease initiation, as APOB-lipoproteins can be retained in the liver by binding to heparan sulphate proteoglycans (HSPG), similar to sub-endothelial retention at the onset of atherosclerosis(13, 14).

Wild-type mice have low-levels of APOB-lipoproteins in circulation, even when subjected to western-style diets, and common genetic and diet-based mouse models of NAFLD often lack an elevated APOB-lipoprotein component. These low-levels of APOB-lipoproteins, with cholesterol largely being transported in non-atherogenic high-density lipoprotein, excludes examining the role of NAFLD in atherosclerosis in wild-type mice. Mouse models of familial hypercholesterolemia such as the apolipoprotein E (*Apoe*^*-/-*^) and LDL receptor deficient (*Ldlr*^*-/-*^) strains, which have been used for decades in atherosclerosis research (15), have recently proven their utility in understanding NAFLD (16). In particular, an inflammatory role for the oxidation-specific epitopes that can form in lipoproteins has been shown in established and late stage NAFLD (9, 17).

The initiation of NAFLD, and specifically the contribution of APOB-lipoproteins to the onset of primary steatosis, remains largely unexplored as the *Apoe*^*-/-*^ and *Ldlr*^*-/-*^ strains are born with elevated plasma cholesterol and hence in a state of allostasis. It is currently unknown which cells in the liver, if any, first respond to the ‘dyslipidemic insult’ of high concentrations of circulating atherogenic APOB-lipoproteins, and if such putative responses are protective or pathogenic towards ACVD. The kinetics of any such response to atherogenic dyslipidemia are also obscure: are days, months or weeks needed before the initiation of liver pathology can be detected?

We hypothesized that rapidly switching the adult mouse into a state of atherogenic dyslipidemia would capture the initiation of liver steatosis mediated by APOB-lipoproteins. Here, we introduce two complementary approaches to achieve hypercholesterolemia, with either inducible deletion of *Apoe* or overexpression of the human PCSK9 D374Y mutation. Liver lipid accumulation was rapid, within 10 days of initiating dyslipidemia, and was accompanied by both common and strain unique liver responses. Subjecting the mice to a high-fat diet revealed an immune dominated response conserved between strains and that correlated with human liver PCSK9 levels. By constructing a reporter mouse strain that allows for *ex vivo* monitoring of cellular APOB-lipoprotein uptake and combining this with single cell RNA-seq (scRNA-seq) and deletion experiments, we reveal that Kupffer cells dominate the liver response through secretion of pro-atherogenic factors but also by restraining circulating APOB-lipoprotein levels. Together, we propose that our understanding of atherosclerosis initiation should expand beyond vasculature-centric models.

## Results

### Mouse models for acute inducible dyslipidemia

We have constructed two mouse models of inducible dyslipidemia through targeting APOE and LDLR. The rationale behind this is to discover common *in vivo* responses following acute dyslipidemia, rather than specific APOE or LDLR gene functions.

We complemented our previously described inducible model of dyslipidemia based upon conditional loss of *Apoe* (18), we have created a second model by inserting the human PSCK9 variant D374Y into the *ROSA26* locus. This was crossed with the tamoxifen inducible and ubiquitously expressed *Cre* line *ROSA26*^*CreERt2*^. Hence both conditional loss of *Apoe* (*Apoe*^*fl/-*^ *ROSA26*^*CreERt2/+*^ herein referred to as APOE cKO) or inducible expression of hPCSK9 D374Y (*ROSA26*^*CreERt2/hPCSK9D374Y*^ herein referred to as D374Y) can be achieved by tamoxifen administration (**FigS1a**).

Plasma hPCSK9 increased 11-fold in D347Y mice versus littermate controls (mean 2325 vs. 213 ng/ml) three days post-tamoxifen (**FigS1b**,**c**) with plasma cholesterol levels already significantly increased at 24hours after tamoxifen dosing (**FigS1d**). We next compared the plasma lipids in these two strains at day 10 after tamoxifen administration on a chow diet (**Fig1a**). Cholesterol levels were increased in both APOE cKO (158 vs 55mg/dl) and D374Y (150 vs 67mg/dl), as were phospholipid levels (89 vs 63mg/dl in APOE cKO and 106 vs 71 mg/dl in D374Y and controls). Noticeably, the D374Y strain displayed increased plasma levels of triglycerides (182 vs 119mg/dl) and glycerol (4.6 vs 3.7mg/dl), which was not observed in the APOE cKO mice (**Fig1b**), whilst free fatty acid levels were comparable. In both the APOE cKO and D374Y strains, cholesterol was incorporated in the VLDL/chylomicron remnant fraction of lipoproteins, whereas in D374Y a more prominent accumulation in the LDL fraction was additionally present as expected (**Fig1b)**. The increase in triglycerides present in D374Y mice were incorporated into the VLDL/chylomicron remnant fraction.

**Figure 1.**
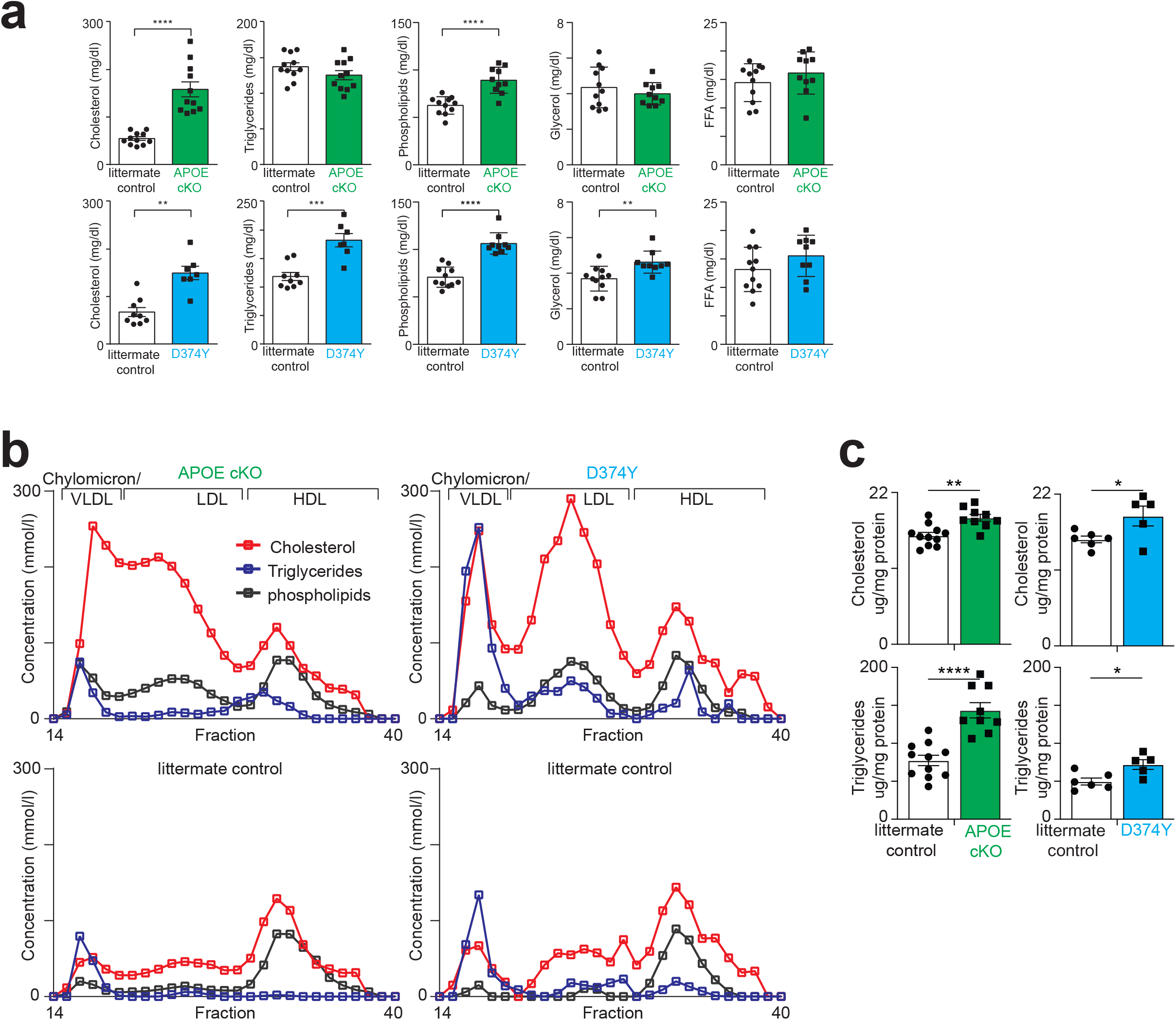
Development of mouse models for inducible steatosis in the adult liver. **a**, Plasma lipid levels (mg/dl) 10 days after tamoxifen administration in APOE cKO (green bars) and D374Y (blue bars) strains together with respective littermate controls in male and female mice fed normal chow. APOE cKO and littermate controls (n = 10-11). D374Y and littermate control (n = 7-11). **b**. Plasma lipoprotein fractionation profiles (µmol/l) at 10 days after tamoxifen dosing. All curves calculated as an average of two separately run plasma pools from male and female mice (plasma from 4-6 mice in each pool). **c**, Cholesterol and triglyceride measurements (µg per mg protein) after Liver Folch extraction from male and female mice APOE cKO and littermate control (n = 9-11) and D374Y and littermate control (n = 9-11). All univariate scatter plots are ± SEM. *p < 0.05, **p < 0.01, ***p < 0.001 and ****p < 0.0001 by t-test

Histological analysis indicated rapid lipid accumulation in the liver (**FigS1e)**. We quantified steatosis initiation using Folch analysis which revealed significant increases in liver cholesterol and triglyceride accumulation in both strains relative to controls already 10 days after tamoxifen treatment and maintained on a chow diet (**Fig1c)**.

In summary, the acutely induced dyslipidemia in adult APOE cKO and D374Y mice resembles that of legacy *Apoe*^*-/-*^ and *Ldlr*^*-/-*^ strains respectively and are suitable for studying the temporal contribution of APOB-lipoproteins to the initiation of NAFLD.

### The liver transcriptional response to HSPG-retained lipoproteins

We next determined how APOB-lipoprotein accumulation affects liver gene transcription. The APOE cKO and D374Y differed considerably in their initial response to acute atherogenic dyslipidemia, with 35 genes differentially expressed in the APOE cKO and 1111 genes in the D374Y (FDR 0.1) as determined by bulk mRNA-seq of the whole liver (**Fig2a**,**b**). A core set of 10 genes were similarly dysregulated in both strains including several known liver macrophage and Kupffer cell identity genes (*Socs2, Folr2, Cd5l, Grn, C6, Il18bp* **Fig2c**,**d**). Most of these 10 genes were also enriched in human Kupffer cells based on data from the human protein atlas (19, 20) (**FigS2a**) APOE cKO showed additional upregulation of myeloid genes (*Ccl24, Cfp, Clec4f, Mpeg1*), virus immunity genes (*Oas2, Oas3, Oasl2, Trex1*), and dysregulation of metabolic factors (*Abcg1, Fabp5, Nr1d1, Scd1, Slc10a2*) including *Apoe* itself (**FigS2b**), whilst unique gene expression signatures in D374Y were dominated by metabolic pathways (**FigS2c**).

**Figure 2.**
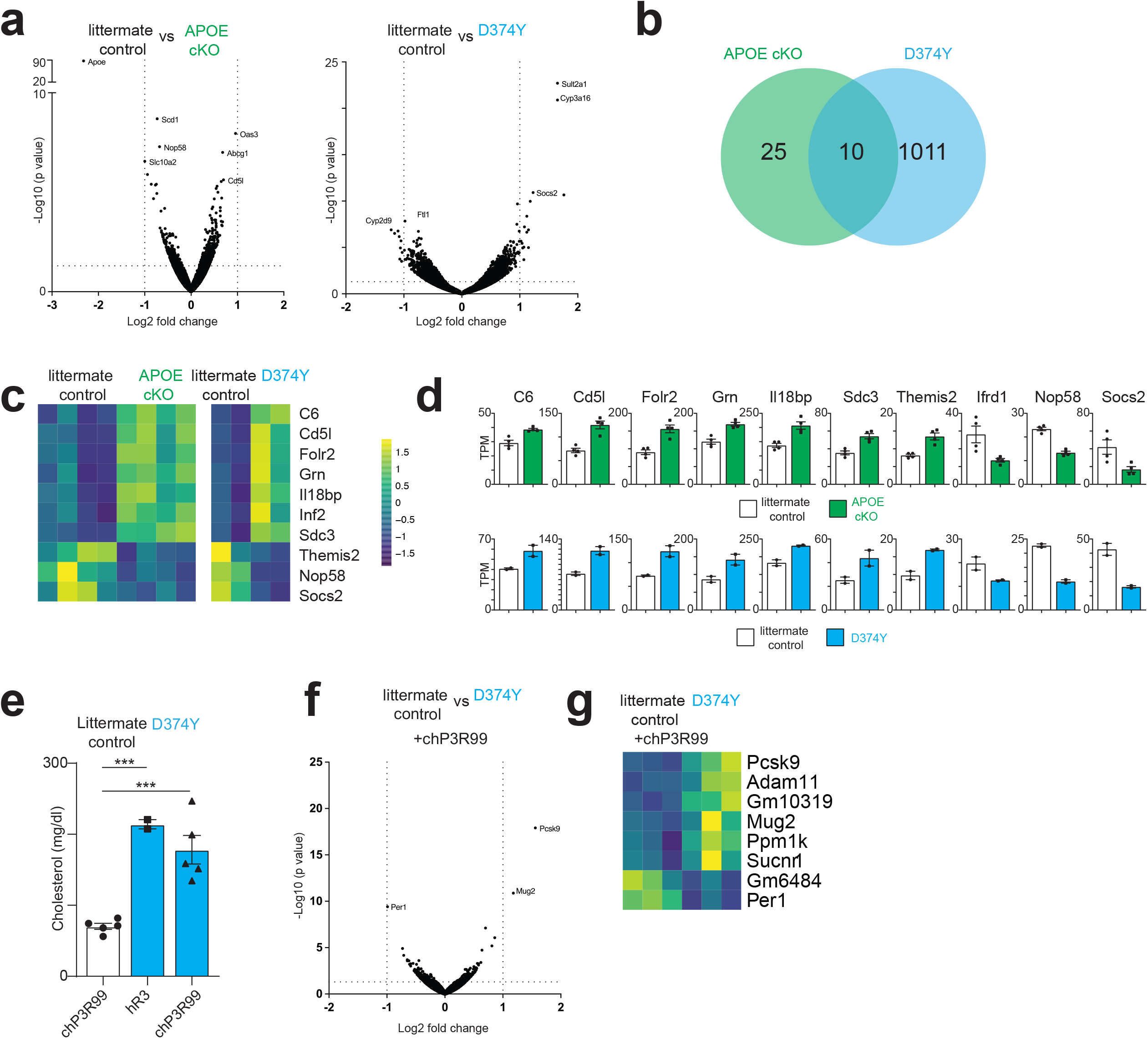
The transcriptional response of the liver to acute dyslipidemia. **a**, Volcano plots of bulk mRNA-seq of whole liver from female APOE cKO (n = 4 vs 4) and D374Y (n = 2 vs 2) mice versus their respective littermate controls. **b**, Venn diagram indicating unique and overlapping differentially expressed genes. **c**, Heatmap of the conserved genes for the two strains. **d**, Transcript per million (TPM) of the conserved genes from each individual liver sequenced. **e**, Cholesterol measurement (mg/dl) of female D374Y and littermate controls given hR3 control or HSPG-binding or chP3R99 antibody. ***p < 0.001 by one-way ANOVA. n=2-5 **f**, Volcano plot from mRNA-seq of female D374Y versus littermate control administered chP3R99 antibody (n = 3 vs 3). **g**, Heatmap of significant differentially expressed genes from female D374Y versus littermate control administered chP3R99 antibody. All experiments day 10 after tamoxifen treatment and maintained on a chow diet.

We next injected HSPG binding antibodies, that have previously been show to prevent APOB-lipoprotein retention in the vasculature (21), into D374Y mice and induced dyslipidemia (**Fig2e**). We now only observed 8 genes that are mostly liver specific differentially expressed, including *Pcsk9* itself (**Fig2f**,**g**), and the 10 conserved genes where no longer dysregulated (**FigS2d**).

Taken together, a transition to atherogenic dyslipidemia results in large-scale gene expression changes in the liver that is dependent on HSPG retention and distinguished by a conserved myeloid cell response.

### A conserved inflammatory response coalesces upon sustained atherogenic dyslipidemia

*Apoe*^*-/-*^ and *Ldlr*^*-/-*^ strains are established NALFD models (22, 23), characterized by inflammation in response to prolonged administration of western-style diets. We next determined if prolonged APOB-lipoprotein dyslipidemia in our APOE cKO and D374Y initiation models also culminates in a conserved inflammatory response. Bulk mRNA-seq of livers from both strains after 12 and 20 weeks of high fat diet revealed a convergence of the gene expression profile between both strains, with 275 genes being similarly differentially expressed at week 12 (**Fig3a**,**b**) and 196 at week 20 (**Fig3c**,**d)**, with 87 genes conserved between both time points. Notably, a majority of these genes were related to immune function, indicating that the liver responds to sustained APOB-lipoprotein dyslipidemia through an inflammatory response (**Fig3e**,**f)**.

**Figure 3.**
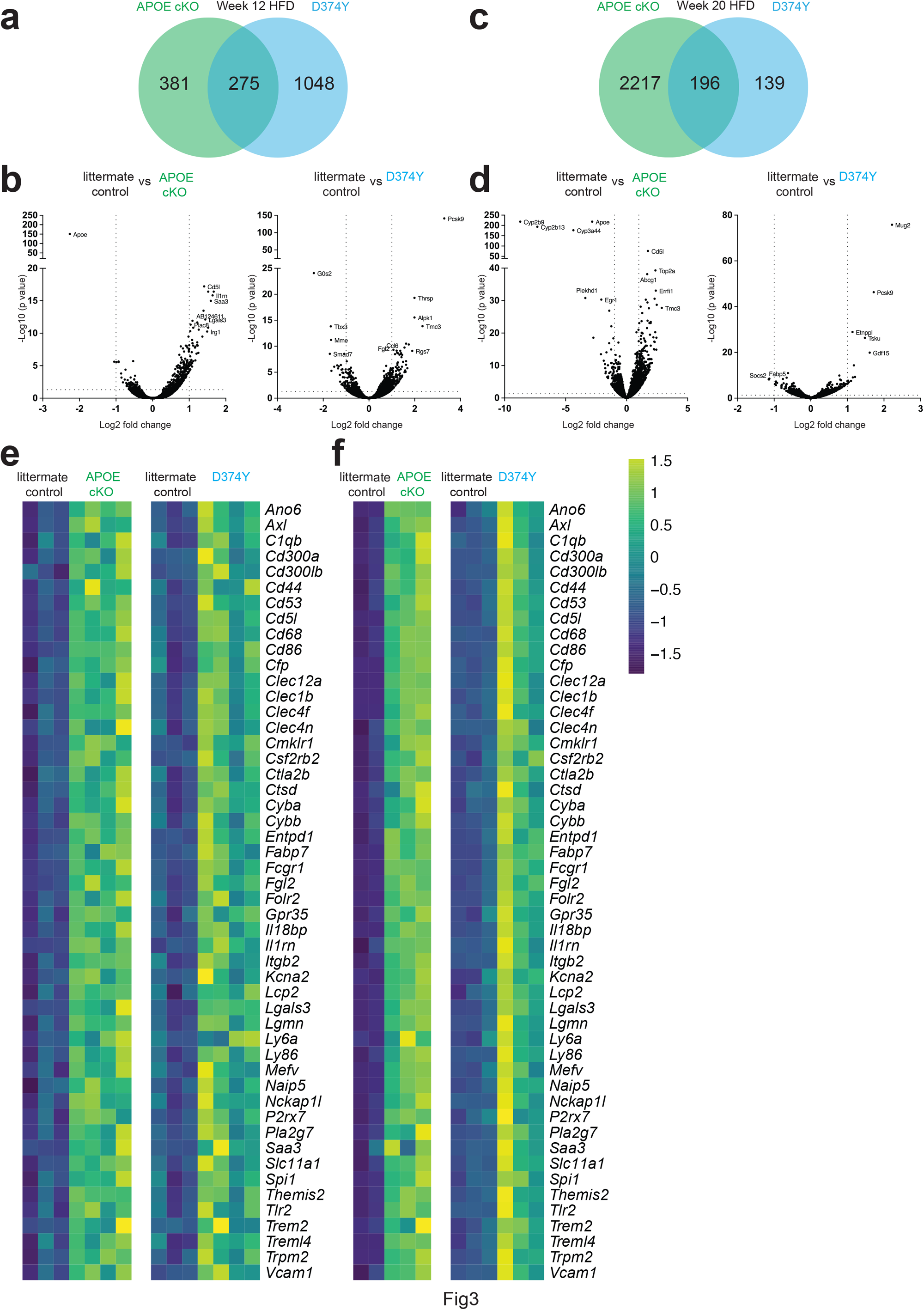
A conserved inflammatory response to sustained dyslipidemia. **a**, Venn diagram indicating differentially expressed genes that are conserved or unique to each strain. **b**, Volcano plot for mRNA-seq of whole liver for male APOE cKO (n = 3 vs 4) and D374Y (n = 3 vs 4) mice and respective littermate controls after 12 weeks of HFD. **c**,**d**, Volcano plot and Venn diagram for mRNA-seq of whole liver for female APOE cKO (n = 2 vs 3) and male D374Y (n = 3 vs 3) mice and respective littermate controls after 20 weeks of HFD. **e**,**f**, Heatmap for selected inflammatory genes at 12 and 20w after tamoxifen administration and maintained on a HFD.

These experiments also allowed us the opportunity to explore how the APOE cKO and D374Y strains individually differed, by examining gene expression changes conserved at week 12 and 20 only within each strain. This revealed that D374Y mice were marked by changes in lipid metabolism whilst the APOE cKO mice had additional dysregulation of immune genes (**FigS3a**,**b**). We next analyzed the 87 conserved genes in human liver samples (n=261), and 42 of these genes also showed significant positive correlation with PCSK9 transcription (**FigS3c**,**d)**.

Altogether, we observed a conserved inflammatory response in the liver to long-term atherogenic dyslipidemia, therefore validating our models as suitable for studying the initiation of liver steatosis.

### Identification of Liver immune cells that recognize APOB-lipoproteins

To characterize which cells were capable of taking up APOB-lipoproteins, we tagged the Apolipoprotein B protein at the N-terminus with the mCherry fluorescent protein (**FigS4a)**, and crossed this together with D374Y strain. Widespread mCherry could be detected in the liver (**FigS4b)**, and cholesterol levels were noticeably increased following tamoxifen administration, and hence APOB function was maintained (**FigS4c)**.

Analysis of the liver by flow cytometry, 10 days after tamoxifen administration and maintained on a chow diet, revealed multiple classes of immune cell positive for mCherry compared to D374Y mice alone (**Fig4a**). We sorted live CD45+ve mCherry cells from the liver and performed 10X single cell RNA sequencing on the cells (**Fig4b**). We identified 10 clusters (**Fig4c**,**S4d)** of cells consisting of cluster 1 (Hepatocytes: *Alb, Mat1a, CPs1*) cluster 2,6 &7 (Kupffer cells: *Adgre1, Timd4, Csf1r*) cluster 3 (Neutrophils: *S100A8, S100A9, Clec4e*), cluster 4 (Macrophages: *Cxc3r1, MHCII, Apoe*), cluster 5 (Endothelial: *Clec4g, Igfbp7,Kdr*), cluster 8 (Dendritic cells: *Cst3, Crip1, Flt3, MHC II*), cluster 9 (T cell: *Trbc2, Cd3g, Lck*) and Cluster 10 (B cell: *Cd79a, Ebf1, Pax5*).

**Figure 4.**
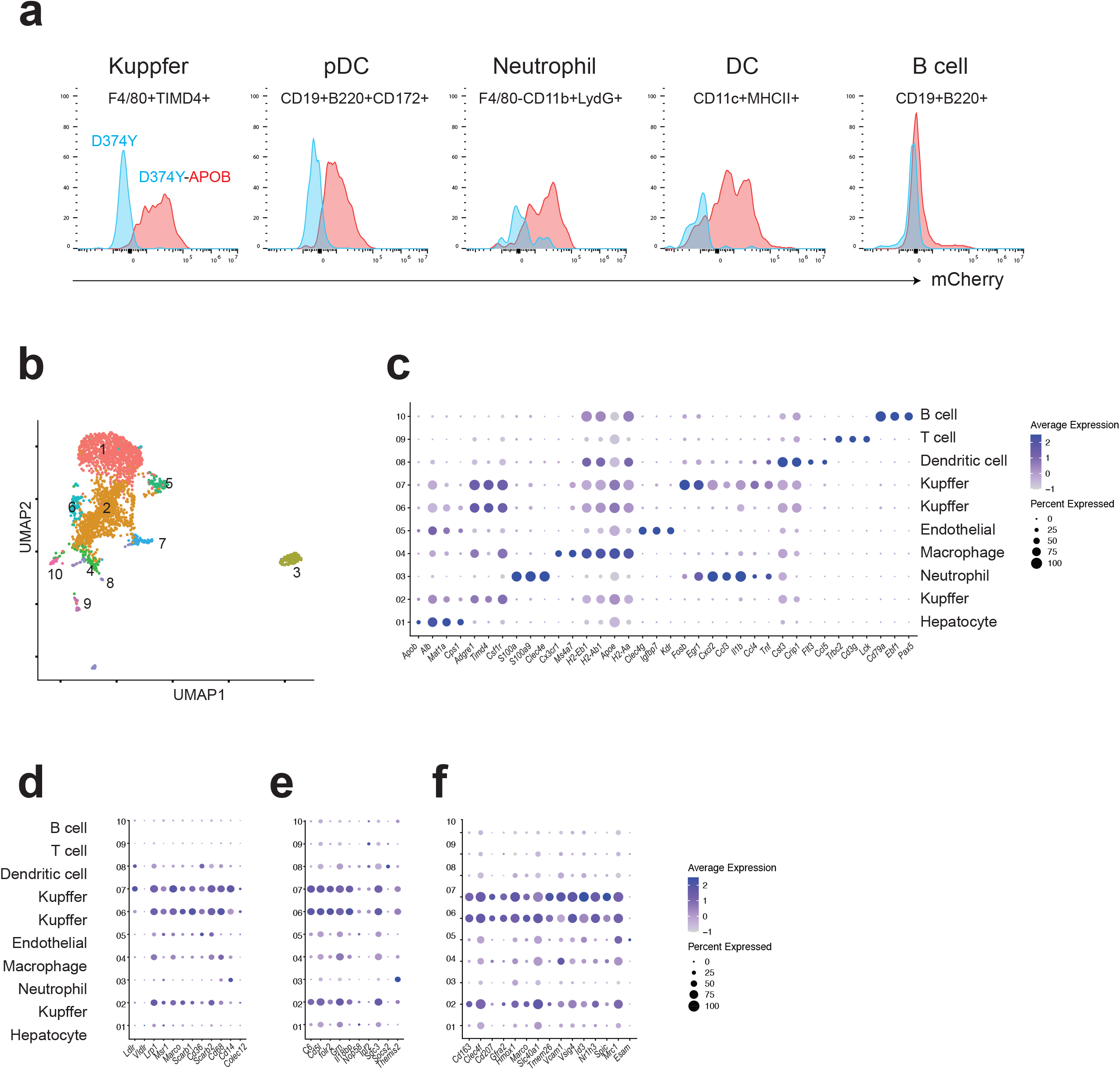
Monitoring uptake of atherogenic lipoproteins through in vivo labelling of APOB-lipoproteins. **a**, Flow cytometry from immune cells extracted from the liver revealing mCherry expression in male D374Y mCher-ry-APOB (D374Y-APOB) or D374Y mice alone. Cells were gated as indicated above each histogram **b**, UMAP plot indicating clusters from scRNA-seq of liver female CD45+mCherry+ cells. **c**, Dot plot for identity markers from clusters 1-10. **d**, Expression of receptors capable of LDL uptake. **e**, Identification of cell clusters expressing the day 10 conserved genes. **f**, Expression of core Kupffer cell identity genes within each cluster. All experiments from mice days 10 after tamoxifen treatment and maintained on a normal chow diet.

Notably the Kupffer cell cluster 7 population additionally expressed of an array of acute inflammatory genes (*Il1b, Tnf, Cxcl12, Ccl3, Ccl4, Fosb, Egr1*). The 3 Kupffer cell populations were additionally enriched for receptors that are known to uptake APOB-lipoproteins (**Fig4d**), and for the conserved set of 10 genes dysregulated at day 10 post-tamoxifen in both APOE cKO and D374Y(**Fig4e**). Additionally, the KCs were positive for the recently identified core set of KC genes (*Cd163, Clec4f, Cd207, Gfra2, Hmox1, Marco, Slc40a1, Tmem26, Vcam1* & *Vsig4*) (24, 25) including those that control lipid-handling (*Id3, Nr1h3* & *Spic*) (26, 27), as well *Mrc1* but not *Esam*. (**Fig4f**). The absolute number of Kupffer cells did not change in the liver (**FigS4e)**.

In conclusion upon transition to a steatotic liver, Kupffer cells can take up a APOB-lipoproteins and induce an inflammatory gene expression profile.

### Liver Kupffer cells coordinate the liver response to APOB-lipoprotein dyslipidemia

To determine the role of Kupffer cells upon the transition to steatosis, we ablated these cells using clodronate liposomes (28) (**FigS5a**) and analyzed the liver 10 days after tamoxifen administration with mRNA-seq. As expected, analysis of the D374Y strain administered with clodronate liposome versus D374Y mice injected with control Dil liposomes, revealed strong downregulation of the Kupffer cell gene expression program such as *Clec4f, Timd4 and Cd5l* (**Fig5a**,**b**). Additionally, we also observed a loss of expression of *Il1b, Ccl5, Ccl6, Ccl24* and *Il6*, indicating these pro-inflammatory cytokines and chemokines are normally expressed in the liver upon transition to acute dyslipidemia. Blood monocytes were not affected by clodronate treatment (**FigS5b**).

**Figure 5.**
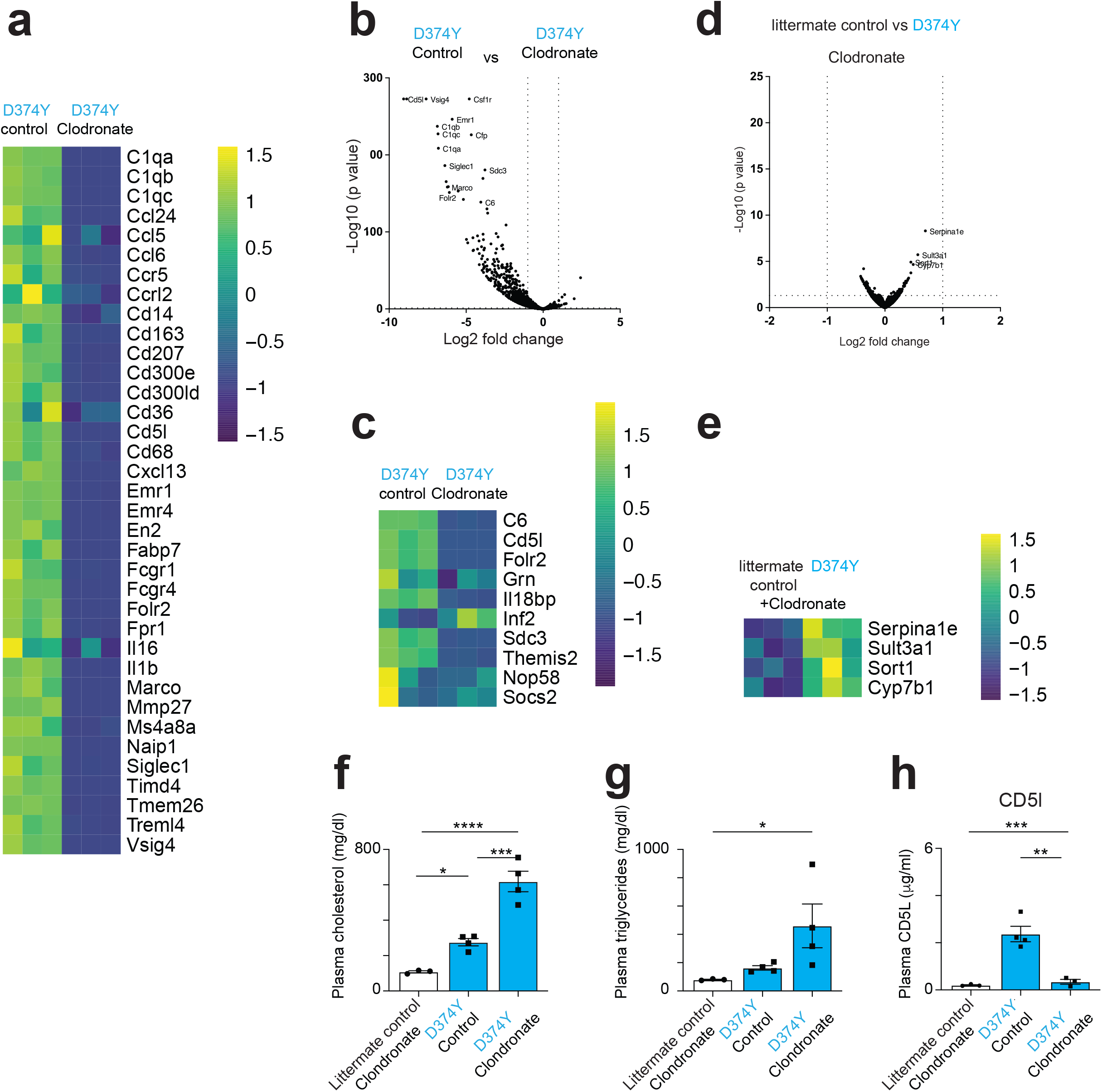
Ablating Kupffer cells prevents the hepatic response to atherogenic dyslipidemia. **a**, Heatmap and **b**, volcano plot of genes downregulated in clodronate liposome versus Dil liposome treated female D374Y mice as determined by mRNA-seq (n = 3 vs 3). **c**, effect of clodronate liposomes on the day 10 conserved gene expression. **d**,**e** Volcano plot and heat map indicating minimal response of the liver to dyslipidemia when comparing littermate control versus D374Y with both treated with clodronate liposomes, as determined by mRNA-seq (n = 3 vs 3). **f**, Total plasma cholesterol and g, triglyceride measurements. **h**, Plasma CD5l concentrations as determined by ELISA *p < 0.05 **p < 0.01 ***p < 0.001 and ****p < 0.0001 by one-way ANOVA. All experiments day 10 after tamoxifen treatment and maintained on a normal chow diet. All univariate scatter plots are ± SEM.

The 10 genes that showed conserved dysregulation in APOE cKO or D374Y at day 10 were no longer differentially expressed, consistent with these being Kupffer cell genes (**Fig5c**). We also compared littermate control mice and D374Y mice, both treated with clodronate liposomes. Strikingly, the large scale gene expression changes normally seen at day 10 in D374Y mice were absent, with only 4 hepatocyte-specific genes (*Serpina1e, Sult3a1, Sort1 and Cyb7b1*) being differentially expressed (**Fig5d**,**e**).

To exclude the possibility that Kupffer cell ablation prevented dyslipidemia as an alternative explanation for this lack of liver response, we measured plasma lipids in our cohort. Loss of Kupffer cells strongly increased both the concentrations of cholesterol (approximately 600mg/dl) and triglycerides (approximately 400mg/dl) compared to controls, even though the mice were maintained on a normal chow diet (**Fig5f**,**g**).

CD5L is a secreted pro-atherosclerotic factor (29) and canonical marker of KCs (27). *Cd5l* was the only factor upregulated in both strains at all time points tested - Day 10 chow diet and week 4, 12 & 20 HFD. ELISA measurement of plasma at day 10 revealed a 10-fold increase in CD5L in Dil liposome-control treated D374Y mice versus clodronate treated littermate controls. Hence dyslipidemic mice versus non-dyslipidemic lacking liver KCs. Clodronate treatment of D374Y mice reversed this increase in CD5L in response to dyslipidemia (**Fig5h**).

In summary, Kupffer cells are essential for the inflammatory liver response to dyslipidemia and restrain plasma atherogenic lipoprotein concentrations.

### The inflammatory response to dyslipidemia requires APOB-lipoproteins and CD8 T cells

To examine the role of Kupffer cells and APOB-lipoproteins during the establishment of liver steatosis, we subjected our mouse strains to a short 4-week high-fat diet (HFD) following Tamoxifen administration. Cholesterol was further increased with this diet and time point compared to day 10 chow **(FigsS6a)** and steatosis and ballooning was clearly visible in both genotypes **(FigsS6b)**. 28 genes were upregulated and 4 downregulated in both APOE cKO and D374Y strains compared to their respective littermate controls as determined by bulk mRNA-seq **(Fig6a,b, FigsS6c)**. Noticeably, those transcripts increased included well-established pro-inflammatory mediators including *Ccl2, Ccl6, Ccl24, Cd5l, Irg1,Lgals3, Lgmn, P2rx7, Pla2g7, Scimp, Sell, Slc15a3 & Tyrobp* and acute phase proteins *Saa1,Saa2 & Saa3*. Analysis of the day 10 scRNA-seq data revealed strong enrichment among the KC clusters for this inflammatory 4-week HFD signature **(Fig6c)**, and many of these genes were again enriched in human KCs based on data from the human protein atlas **(FigsS6d)**. Circulating CD5L was also increased both in week 4 HFD APOE cKO and D374Y CD5L plasma **(Fig6d)**.

**Figure 6.**
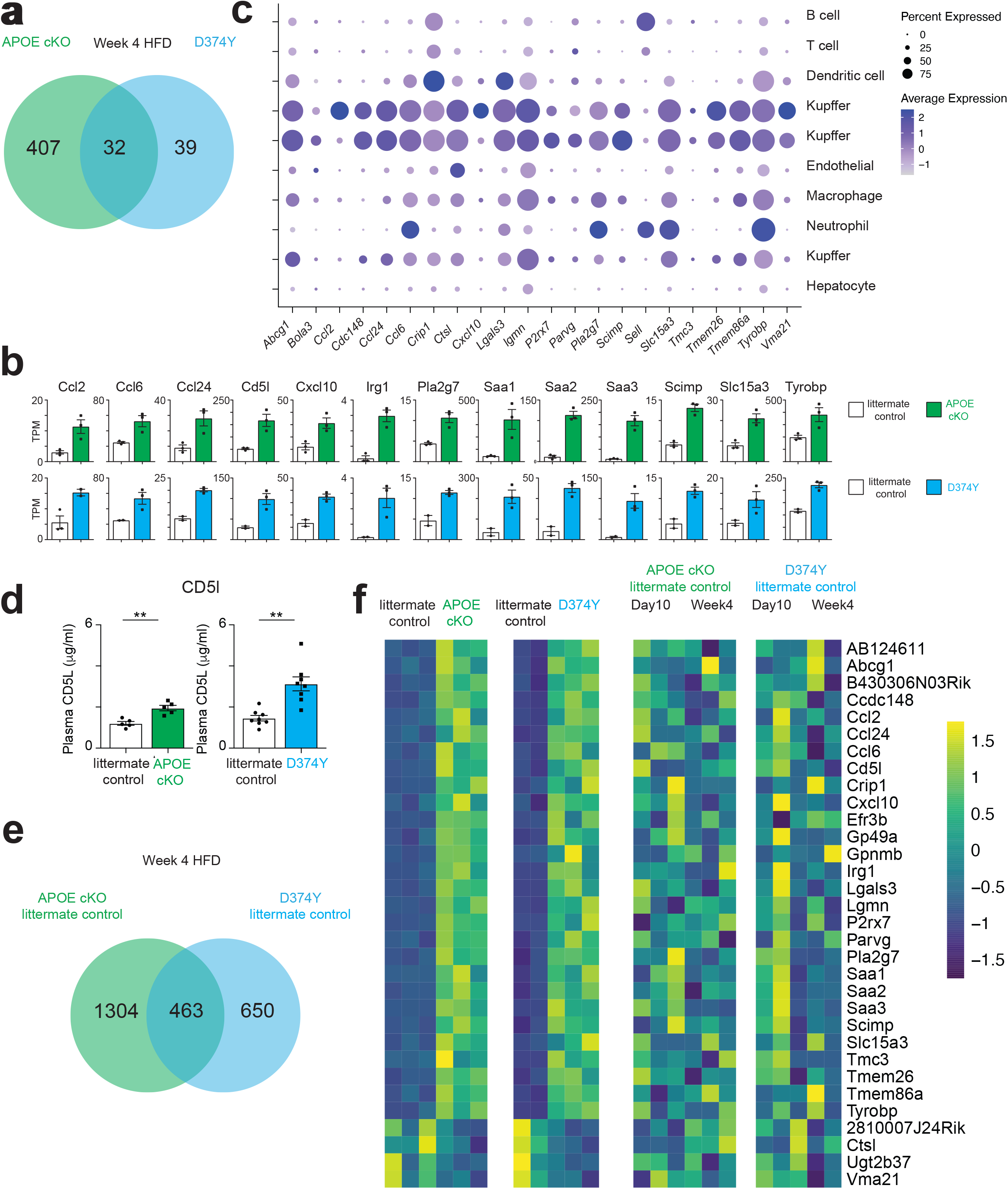
APOB-lipoproteins are required for the inflammatory response to a high-fat diet. **a**, Venn diagram of unique and overlapping differentially expressed liver genes after 4 weeks of high-fat diet following tamoxifen-induction in female APOE cKO (n = 3 vs 3) and D374Y mice (n = 2 vs 3) versus respective littermate controls, as determined by bulk mRNA-seq. **b**, TPM values for selected conserved genes from both strains. **c**, Expression of these conserved genes from this week 4 HFD analysis in the liver day 10 scRNA-seq analysis. **d**, ELISA for secreted CD5L in plasma (APOE cKO n= 5 vs 5, D374Y n=8 vs) *p < 0.01 by t-test. **e**, Identification of conserved transcriptional response to 4 weeks of HFD from littermate controls only of both strains. **f**, Heatmap for expression of conserved genes from APOE cKO and D374Y mice compared to littermate controls alone in response to 4w of HFD.

Next using only the littermate controls we compared the gene expression changes between day 10 chow-fed and week 4 HFD-fed mice, and hence largely in the absence of atherogenic APOB-lipoproteins, to determine the effects of HFD alone on the liver. We identified strong overlaps between the littermate controls of both strains, with 463 genes similarly dysregulated **(Fig6e)**, with metabolic pathways enriched **(FigS6e)**. Notably however, none of the 32 genes differentially expressed after 4 weeks of HFD in the APOE cKO and D374Y strains were dysregulated by high-fat diet alone, indicating that this specific pro-inflammatory response requires APOB-lipoproteins **(Fig6f)**.

CD8 T cells are required for the maintenance of adipose macrophages (30) and liver Kupffer cells (31). We antibody-ablated CD8 T cells during this 4-week HFD regime **(FigS7a)** and observed a down-regulation of the Kupffer cell gene expression program in APOE cKO and D374Y strains (**Fig7a**,**b**, **FigS7b**). 22 of the 24 conserved and differentially regulated genes from both APOE cKO and D374Y following CD8 depletion overlapped with those that were down-regulated following clodronate administration at day 10 **(FigS7c)**. Similar to the clodronate administration experiment, cholesterol and triglyceride levels were also increased upon loss of CD8 T cells **(Fig7c)**. None of the 28 genes upregulated in both strains after 4-week high-fat diet were still upregulated when both strains were treated with anti-CD8 **(Fig7d)**.

**Figure 7.**
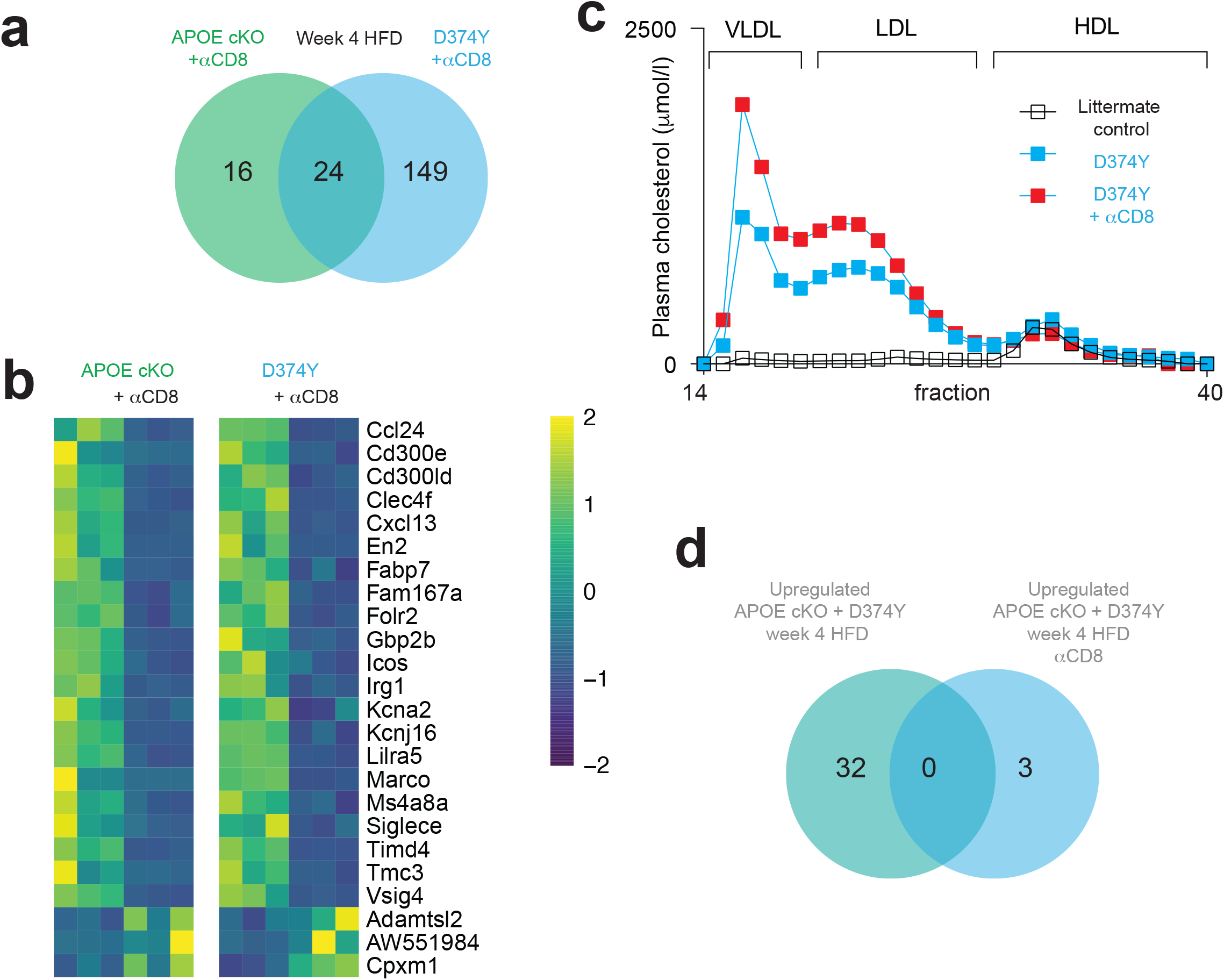
CD8 T cells maintain Kupffer cell responses to atherogenic dyslipidemia. **a**, Venn diagram representing differentially regulated genes in female APOE cKO and D374Y mice, 4 weeks of high-fat diet following tamoxifen-induction, and treated with anti-CD8 antibody relative to PBS treated mice as determined by mRNA-seq. **b**, Heat map of conserved genes differentially expressed in both strains following anti-CD8 treatment. **c**, Plasma lipoprotein fractionation profiles (µmol/l). All curves calculated as an average of plasma pooled from female mice (n=5-6). **d**, Overlap of genes upregulated in both APOE cKO and D374Y mice versus those upregulated in both APOE cKO and D374Y mice but treated with anti-CD8 antibody, in all cases after 4 weeks of high-fat diet following tamoxifen-induction.

Finally as an additional control, we deleted B cells in D374Y mice using anti-CD20 (**FigS7d)**, as these can also uptake mCherry-APOB, but are not known to deplete KCs. In contrast to anti CD8 treatment, no loss of the KC gene expression program was now observed (**FigS7e)**.

## Discussion

APOB-lipoproteins have a decisive pathological function in the development of atherosclerosis and can also contribute to NAFLD when concentrations are raised above physiologically required concentrations (1, 3). The existence of responses outside of the vasculature to the abrupt onset of an atherogenic lipid profile has remained unknown. Here, we have generated mouse models to investigate how and where APOB-lipoprotein retention promotes tissue damage based on inducible loss or gain of function alleles that mimic human familial hypercholesterolemia. Our experimental approach takes advantage of the fact that we can abruptly switch adult mice into atherogenic lipid profile, and hence have been previously untouched by this dyslipidemic insult.

Liver Kupffer cells rapidly responded and functioned as key liver cells responsible for signaling a systemic response at the onset of atherogenic dyslipidemia. This conclusion is based on multiple lines of evidence. Firstly, bulk mRNA-seq analysis of the liver revealed know KC genes being differentially regulated upon transition to dyslipidemia. Secondly, through use of the mCherry-APOB reporter mouse, KCs were shown to recognize APOB-lipoproteins in the context of atherogenic dyslipidemia. KCs also dominated the landscape of an unbiased scRNA-seq analysis of mCherry-APOB positive immune cells. Deleting KCs directly through clodronate liposomes almost completely abolished the whole liver transcriptional response to acute dyslipidemia in the D374Y strain. Indirect deletion of KCs through anti-CD8 administration also extinguished the KC-specific inflammatory response whereas control anti-CD20 did not. KCs also expressed multiple receptors that are known to facilitate APOB-lipoprotein uptake.

One subset of KCs was enriched for proinflammatory mediators such as *Il1b, Tnf and Ccl4*. One possibility that requires further experimentation is that this inflammatory subset represents a differentiation step occurring after APOB-lipoprotein uptake. KCs have recently been subdivided in KC1 and KC2 populations, with KC2 reported to regulate metabolism (32, 33), although this interpretation has been challenged (34). We only observed *Esam* expression, used to define KC2, in the endothelial cluster. Hence APOB-lipoprotein responses by KCs doesn’t seem to involve the KC2 subset.

Notably, deletion of KCs dramatically increased plasma cholesterol and triglyceride levels in D374Y mice. KCs in rabbits, rats and humans are known to take up considerable amounts of circulating LDL (4-6). Taken together with observations from *Apoe*^*-/-*^ mice in the specific context of high fat diet and hereditary haemochromatosis (7), our results further underscores the critical function of KCs in regulating plasma lipoproteins levels. Therefore, KCs have a potentially ambivalent function, by expressing and secreting pro-inflammatory factors but at the same time reducing circulating pro-atherogenic APOB-lipoproteins. In this sense KCs function as hepatic sentinels of atherosclerosis initiation.

The response to retention hypothesis of atherosclerosis proposes that sub-endothelium retention of APOB-lipoproteins within the vasculature is the key initiating step in plaque development (3). Perhaps less well appreciated is the existence of a similar process in the liver (35). In our own preliminary analysis, APOB could be visualized retained at susceptible sites in the vasculature already 10 days after the initiation of dyslipidemia, resulting in large transcriptional changes in both APOE cKO and D374Y (36). However, in contrast to the liver phenotype reported here, this aortic response showed little conservation between the two strains. A provocative interpretation is that the liver, particularly KCs, have a central role in responding to the initiation of atherogenic dyslipidemia, beyond or preceding that of the vasculature. Hepatic responses may further modulate plaque responses at distant sites. In line with this, we observed increases at both the transcriptional level (*Ccl2, Cd5l, Saa2/3, Pla2g7*) and with systemic levels of secreted proteins (CD5L) with factors know to alter atherosclerosis burden. This raises the question as to what the long-term effects of KCs are on atherosclerosis development. Such experiments will be difficult to address, as KCs are needed for mouse viability over the long-term. However, individually targeting these factors may provide insight. One possibility is that there are multiple subsets of liver macrophages in response to raised levels of APOB-lipoproteins, similar to aortic phagocytic cells (37).

Here, we have shown that transitioning to a state of acute atherogenic-dyslipidemia results in large-scale transcriptional changes in the liver upon accumulation of cholesterol and triglycerides. Multiple plausible processes could explain this lipid accumulation; uptake of circulating APOB-lipoproteins which are increased in concentration due to the defect in hepatic lipoprotein clearance in our mouse strains. Secondly, it could be due to an increase in *de novo* lipogenesis or defects in Beta-oxidation, or finally it could be a result of inefficient lipoprotein secretion by hepatocytes (38). Administering a specific antibody that has established efficacy in preventing APOB-lipoprotein retention prevented liver transcriptional responses upon steatosis. This would indicate that the liver transcriptional response to acute dyslipidemia is dependent on retained extracellular APOB-lipoproteins. However, further studies will be needed to investigate the role of hepatic *de novo* VLDL lipogenesis and secretion upon the onset of acute atherogenic dyslipidemia.

Subjecting our mice to a sustained high-fat diet over many weeks revealed considerable convergence in the whole liver transcriptional response in both strains, dominated by genes associated with immune function, underlining the inflammatory nature of chronic steatosis. This prominent inflammatory signal is specific to hypercholesterolemic APOE cKO and D374Y strains, as litter-mate controls did not respond to a HFD in a similar manner. These results are consistent with a recent report indicating a non-inflammatory role of macrophages in mouse models of non-alcoholic steatohepatitis that were also lacking an atherogenic dyslipidemic component (39). Noticeably, the long-term HFD treatment resulted in more inflammatory responses in the APOE cKO than the D374Y strain, consistent which an anti-inflammatory role of APOE beyond lipoprotein transport (18, 40).

Approximately half of the total genes that were upregulated in both strains at both 12 and 20 weeks were positively correlated with human liver PCSK9 expression levels. Similarly, the mouse Kupffer cell genes that are dysregulated in response to atherogenic dyslipidemia are also enriched in human Kupffer cells. Our approach of combining two inducible dyslipidemic strains may therefore be suitable as a pre-clinical model for understanding how perturbations in lipoprotein-handling can lead to systemic responses. Indeed, mouse models have proven very useful for the studies of atherosclerosis (41). However the strains described here, as well as legacy *Apoe*^*-/-*^ and *Ldlr*^*-/-*^ mice, remain models of familial hypercholesterolemia and likely represent the far-end of the spectrum of disease states caused by elevated levels of APOB-lipoproteins.

In summary, we have created an experimental platform that allows for the discovery of how atherogenic dyslipidemia can initiate and sustain tissue damage. Utilization of these tools has identified liver Kupffer cells as key mediators of the hepatic response at the onset of atherosclerosis. Further investigation using these strains will likely allow for the discovery of additional organs that have a pathological response to APOB-lipoprotein mediated dyslipidemia.

### Materials and Methods

### Mice and experimental diets

The APOE cKO mice have been described earlier (18, 42) and were maintained on the C57BL/6 genetic background. All mice were housed in specific pathogen free vivarium at the Karolinska institute. The light/dark period was 12h/12h, and all the mice had *ad libitum* access to food and water. Mice in breeding were fed chow diet R36 (12.6 MJ/kg, 18% protein, 4% fat; Lantmännen, Sweden). Experimental mice received chow diet (R70, Lantmännen, Sweden, 12.5 MJ/kg, 14% protein, 4.5% fat) or HFD (R638, Lantmännen, Sweden, 15.6 MJ/kg, 17.2% protein, 21% fat, 0.15% cholesterol) as stated in each experiment. The Stockholm board for animal ethics approved the experimental protocols.

### Generation of hPCSK9 D374Y and mCherry-APOB mouse strains

We created a conditionally-activated hPCSK9 D374Y gain-of-function mouse model by inserting D374Y mutated hPCSK9 into *Rosa26* locus. The ROSA26 gene-targeting vector was constructed from genomic C57BL/6N mouse strain DNA (GenOway, Lyon, France). PCSK9 D374Y human sequence was inserted downstream of a lox-STOP-lox cassette. When the floxed STOP cassette is removed by CRE recombinase, human PCSK9 D374Y expression is driven by the CAG promoter. The linearized targeting vector was transfected into C57BL/6 embryonic stem cells (ES cells) (GenOway, Lyon, France) according to GenOway’s electroporation procedures. The *ROSA2*^*PCSK9D374Y*^ mice were crossed with *ROSA26*^*CreERt2*^ mice (43), creating *ROSA26*^*CreERt2/PCSK9D374Y*^ experimental mice. Littermates without the D374Y insert (*Rosa26*^*CreERt2/+*^ *or Rosa26* ^*CreERt2/ CreERt2*^) were always used as controls.

For the Knock-in insertion of mCherry into exon 2 downstream of the *Apob* signal peptide in exon 1, a flexible linker ((GGGGS)x3) was inserted between the mCherry sequence and the *Apob* sequence encoding for its mature form. G-418 resistant ES cell clones were subsequently validated by PCR, using primers hybridizing within and outside the targeted locus, and Southern blot, to assess the proper recombination event on both sides of the targeted locus. Recombined ES cell clones were microinjected into C57BL/6 blastocysts, and gave rise to male chimeras with a significant ES cell contribution. The *Apob*^*mCherry/+* was^ bred with C57BL/6 Cre-deleter mice to remove the neomycin cassette between exon 2 and 3 to generate heterozygous mice carrying the reporter allele. These *Apob*^*mCherry/+*^ were then bred to homozygosity and further crossed with *ROSA26*^*CreERt2/PCSK9D374Y*^ to create *Apob*^*mCherry/mCherry*^ *ROSA26*^*CreERt2/PCSK9D374Y*^ *and Apob*^*mCherry/mCherry*^ *ROSA26*^*CreERt2/ CreERt2*^ littermate controls.

### Induction of dyslipidemia

Mice were induced with Tamoxifen at the age of 10-14 weeks. Induction of the models with oral dose of tamoxifen was performed as we previously described (18). We administered 9 mg of tamoxifen dissolved in 150µl peanut oil + 10% ethanol, via single-dose oral gavage, to experimental mice and their littermate controls. The induction efficiency was evaluated by measuring the plasma cholesterol levels. Occasional APOE cKO and D374Y mice that did not show the expected elevation in cholesterol levels were excluded from the study.

### Plasma lipid analyses and Enzyme-Linked Immunosorbent Assay (ELISA)

Blood was drawn via cardiac puncture to EDTA-coated tubes and centrifuged at room temperature for 5 minutes at 500 G. Plasma was separated and stored at −80°C for further analyses. The plasma total cholesterol and triglycerides were measured from plasma with enzymatic colorimetric kits (Randox laboratories) according to manufacturer’s instructions.

Phospholipids (PL) were measured with Phospholipids C kit (Fujifilm, Wako Diagnostics). Plasma free (non-esterified) fatty acid concentrations were measured by an enzymatic colorimetric method (NEFA-HR(2), Wako Chemicals). Plasma concentration of glycerol was determined by an enzymatic colorimetric assay (Free glycerol FS, DiaSys, Diagnostic Systems GmbH). Lipoprotein fractionation was performed for plasma pools of four to six mice by fast performance liquid chromatography method (44). The concentration of cholesterol, triglycerides and PL in each fraction was measured by enzymatic methods using CHOD-PAP kit (Roche Diagnostics) GPO-PAP kit (Roche Diagnostics) and Phospholipids C kit (Fujifilm, Wako Diagnostics), respectively.

The concentration of hPCSK9 was measured in tail vein blood by using the Human PCSK9 Quantikine ELISA kit (R&D Systems, Catalog no. DPC900, Minneapolis, USA) according to kit instructions. The lower limit of quantification (LLOQ) of hPCSK9 in mouse plasma was 625 pg/ml. Mouse CD5L was quantified using an ELISA Kit (Invitrogen, Catalog no. EM15RB) with a LLQQ = 8.19 pg/ml

### Quantification of Liver Lipids

Liver lipid extraction according to Folch was performed as previously described in detail (45). Briefly, Liver lipids were extracted by adding 5 mL of Folch solution (chloroform: methanol - 2:1 v/v) to 100 mg of snap-frozen samples. After serial steps of drying including the addition of 1 mL (0.9% NaCl) to separate the phases and addition of 1 mL of 1% Triton X-100 in chloroform, lipids solubilized in 0.5 mL of water were obtained. Concentrations of the cholesterol and triglycerides in the total lipids was determined using CHOL2 and TRIGL kits (Roche diagnostics). CHOL2 has a measuring range of 3.86–800 mg/dL and TRIGL has a measuring range of 8.85–885 mg/dL. We corrected the total lipids by the protein content in liver though using modified Lowry microassay in plate protocol, with the DC™ Protein Assay (Bio-Rad) kit, with a protein concentration range of 5–250 μg/ml.

### Histology and immunofluorescence

Hematoxylin and eosin or Oil Red O staining for of Liver was performed on 5-um-thick, 4% Zn-formaldehyde-fixed, paraffin embedded liver sections. Sections were deparaffinized in Histolab Clear and rehydrated in gradually decreasing ethanol solutions (99%, 95%, and 70%). They were then stained with hematoxyllin (Vector Labs, Bionordiska) and eosin (Histolab), dehydrated in ethanol and xylene, and mounted in Pertex (Histolab). Oil Red O was prepared as previously described (46). Slides were scanned using a VS200 slide scanner (Olympus) and pictures acquired using OlyVIA V3.4.1 software.

For confocal microscopy of the liver, 10 μm thick sections were thaw-mounted to slides which were subsequently fixed with ice-cold acetone for 10 minutes and stored at −20°C. Slides were thawed and sections were incubated in washing buffer (1x Tris-buffered saline with 0.1 % Tween20) for five minutes. Blocking was performed with Avidin-biotin blocking kit (Vector) according to supplier instructions, and with 5% BSA for 30 minutes. Primary antibodies against CD68 (clone MCA1957, Serotec, 1:10000) were incubated at +4°C over night. The secondary antibody (biotinylated anti-goat IgG (Vector), 1:300) was incubated for 1 hour at RT followed by 1 h incubation with streptavidin-DyLight 647 (Vector, 1:300) subsequently 4′,6-diamidino-2-phenylindole (DAPI) (Invitrogen, 1:50000) for 20 minutes. The sections were washed 3 × 3 minutes in washing buffer after each incubation step. The slides were mounted with fluorescent mounting media (Dako). Negative control stainings were performed by omitting the primary antibodies from the protocol. For CD68+ cells, staining in spleen was used as a positive control. Immunofluorescence images were taken with Nikon Ti-2E confocal microscope using NIS Elements software.

### Bulk and single cell RNA sequencing

Mice were euthanized with carbon dioxide and perfused by infusing PBS via left ventricle. A small sample from one liver lobe was taken, carefully avoiding the gall bladder. Livers were stored in RNALater (Qiagen) at −20°C until RNA extraction. The livers were solubilized in Qiazol lysis reagent using Tissuelyser (Qiagen) and extracted RNA was isolated to upper fraction by chloroform. Purification was performed with RNeasy mini kit including on-column DNAse treatment, using a Qiacube robot. RNA was selected using Poly(A) RNA Selection Kit (Lexogen) and sequencing libraries prepared with Lexogen QuantSeq V2. DNA fragments 200-800 bp for RNA-seq were selected. Cluster generation and sequencing was carried out by using the Illumina HiSeqV4 system with a read length of 50 nucleotides (single-read) or NovaSeq with a read length of 150 nucleotides (paired-end) and aligned to the mouse transcriptome (genome assembly version of July 2007 NCBI37/mm9) using TopHat version 1.4.1. Reads per gene was counted using HTseq version 0.5.3 with the overlap resolution mode set to union. Analysis of differential expression of mRNA was using the DeSeq2 software at default settings, with a false discovery rate set at 0.1 for all experiments as originally described (47).

For scRNA-seq, live CD45+ mCherry+ cells were sorted from the liver. A small samples from one lobe was cut into small pieces and digested for 0.2 mg/mL collagenase IX for 30 min at 37C in RPMI 1640. The cell suspension was passed through a 18g syringe 10 times to remove any clumps and then a 70uM cell strainer. The cell suspension was then washed with FACS buffer before being sorted by a Sony SH800S cell sorter. Cells were processed immediately for GEM generation and barcoding on a 10X Chromium using Next GEM 3’ v3.1 reagents (10X Genomics) followed by sequencing and processing on Cellranger. Data analysis was performed in R using Seurat v4 (48).

### Human samples and data

Liver biopsies were obtained from patients undergoing aortic valve and/or ascending aortic surgery as part of the Advanced Study of Aortic Pathology (ASAP) (49). All protocols were approved by the ethics committee of the Karolinska Institutet, and informed consent was obtained from all participants according to the Helsinki Declaration. Gene expression was analyzed in liver samples (n=261) using the Human Transcriptomic Array 2.0 (Affymetrix). All samples were hybridized and scanned at the Karolinska Institute Affymetrix core facility. Cel-files were pre-processed using Robust Multichip Average (RMA) normalization as implemented in the Affymetrix Power Tools 1.10.2 package apt-probeset-summarize. Expression values were log2-transformed as part of the RMA normalization.

The Single Cell Type section of the Human Protein Atlas contains data from human scRNA-seq experiments retrieved from published studies based on healthy human tissues (19, 20). Transcriptomic expression data for 76 consensus single cell types was obtained from the Human Protein Atlas for the two sets of conserved genes between mouse and human. Analyses were performed using R Statistical Software (50) and the package “pheatmap” (51) was used to visualize expression profiles and the dendrograms were obtained by hierarchical clustering of distances based on gene expression levels for all single cell types using Ward’s criterion.

### Detection of cell types by flow cytometry

Spleens were ground with syringe plungers and prepared to single cell suspensions by pressing through sterile 70 µm mesh size cell strainers. Cells were stained with conjugated antibodies on ice for 30 minutes. The following antibody clones were used: CD3ε (PB 500A2), CD8 (FITC 53-6.7), CD4 (BV750 GK1.5), CD45 (V500 30-F11), CD19 (PE eBio1D3), B220 (APC-Cy7 RA3-6B2), CD172 (BV711 P84), Ly6G (PB 1A8), CD11b (PerCP/Cyanine5.5 M1/70), CD11c (APC N418), MHCII (AF700 M5/114.15.2), F480 (BV510 BM8), TIMD4 (PerCP-eF710 54). Immune cell populations were defined as: Liver Kupffer cell (CD3-CD19-Ly6G-F4/80+TIMD4+ CD64+), Liver pDC (CD19-B220+CD172+), Liver neutrophil (F4/80-CD11b+Ly6G+), Liver DC (CD11c+MHCII+), Liver B cell (CD19+B220+), Spleen CD8 T-cell (CD3+CD4-CD8+) and Spleen B cell (CD45+CD19+B220+). The samples were acquired with Cytek Northern Light or Aurora spectral flow cytometers (Cytek) and analyzed with FlowJo software.

### Antibody administration and lymphocyte cell depletion

The chP3R99 mAb that recognizes sulfated glycosaminoglycans, or hR3 control antibody (52) was administered subcutaneously (100µg per mouse) at day 0 and 6 following Tamoxifen induction and mice maintained on normal chow diet.

CD8 cells were depleted with anti-CD8a antibody (Rat IgG2b anti-mouse CD8α YTS 169.4, BioXcell) or anti-mouse CD20 (Mouse Ig2c anti-mouse CD20 MB20-11 BioXcell). Mice were injected 250 µg antibody in 150µl PBS (anti-CD8) or 100µg in 200 μL (anti-CD20). or similar volume PBS intraperitoneally once weekly. Five days after the first injections, tamoxifen was dosed via oral gavage and the mice were switched to receive HFD. The diet and weekly injections continued for four weeks.

### Statistical analysis

Data are presented as means ± SEM Unpaired Student’s t test used for comparing two experimental groups as indicated. One or Two-way ANOVA followed by fisher’s LSD, or repeated measures ANOVA was used for comparisons of more than two groups as indicated. P < 0.05 was considered significant. Statistical analyses were performed using GraphPad Prism software.

## Acknowledgements

This work was supported by grants from the Leducq foundation Networks of Excellence Program B cells in Cardiovascular Disease, European Community’s Seventh Framework Program FP7-2007–2013 under Grant HEALTH-F2-2013-602114 (Athero-B-Cell), Swedish Research Council (project 2020-01593), the Swedish Heart-Lung Foundation (20210532 and 20210520), a private donation from Fredrik Lundberg and the Novo Nordisk-Karolinska Institutet postdoctoral fellow program.

---

**Figure S1.**
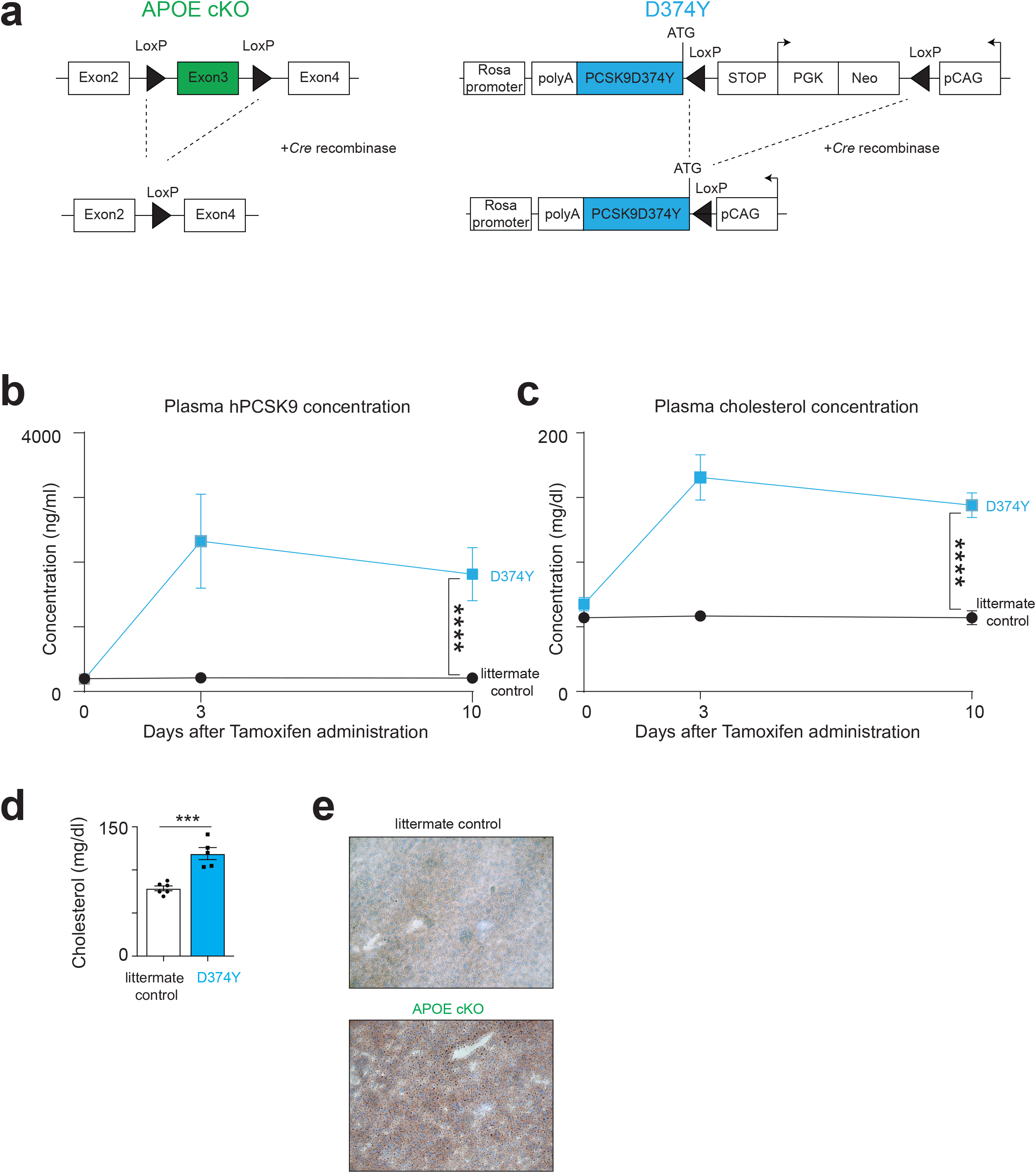
Characterization of mouse models of acute inducible dyslipidemia. **a**, Schematic diagram of the APOE cKO and D374Y alleles. **b**, Enzyme-linked immunosorbent assay for plasma hPCSK9 protein in inducible male and female D374Y mice (blue line) and their littermate controls (black line) before and at 3 and 10 days after tamoxifen dosing. n=2-6. **c**, Plasma total cholesterol measurements in the same time points as in b. n=3-6. **d**, Cholesterol measurements 24 hours after tamoxifen administration from inducible male and female D374Y mice, n=5-6. **e**, Oil red O stained liver section from APOE cKO and littermate control. All plots are mean ± SEM, Statistical analysis was performed with mixed model 2-way ANOVA (b,c) or t-test (d). ***p < 0.001, ****p < 0.0001.

**Figure S2.**
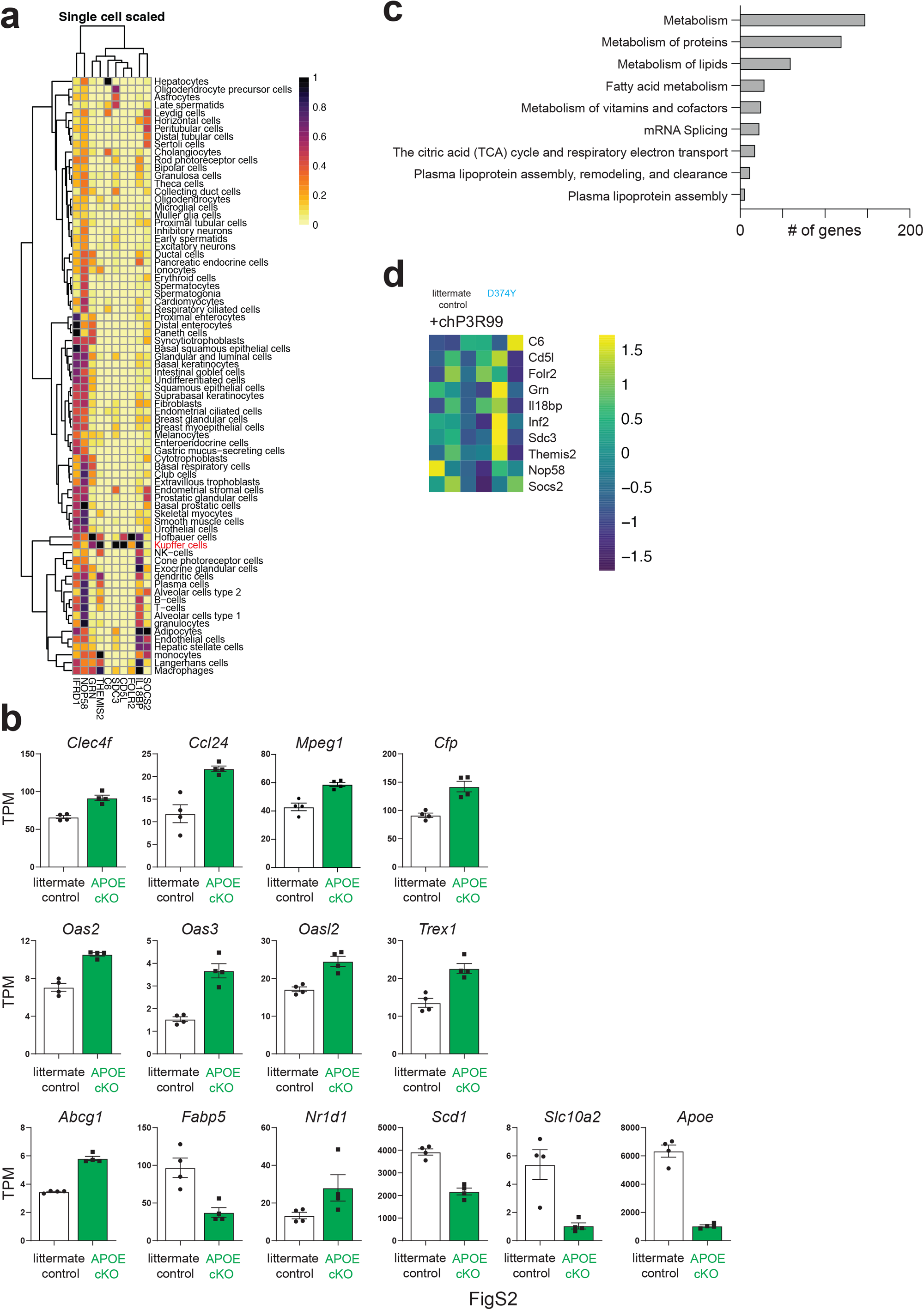
Conserved genes from the mouse are enriched in human Kupffer cells and annotation of pathways unique to either APOE cKO or D374Y strains. **a**, Expression in 76 human single cell types of the 10 conserved genes from APOE cKO and D374Y alleles day 10 after tamoxifen. **b**, TPM (transcript per million) of genes differentially regulated only in the APOE cKO strain All plots are ± SEM and each point represents an individual liver (n = 4 vs 4). **c**, Reactome pathways of genes differentially regulated only in the D374Y strain. Bulk mRNA-seq was performed 10 days after tamoxifen dosing on whole liver in female mice aged 10-12 weeks at induction. **d**, Heatmap for expression of the 10 conserved genes from APOE cKO and D374Y strains in D374Y versus littermate control administered chP3R99 antibody.

**Figure S3.**
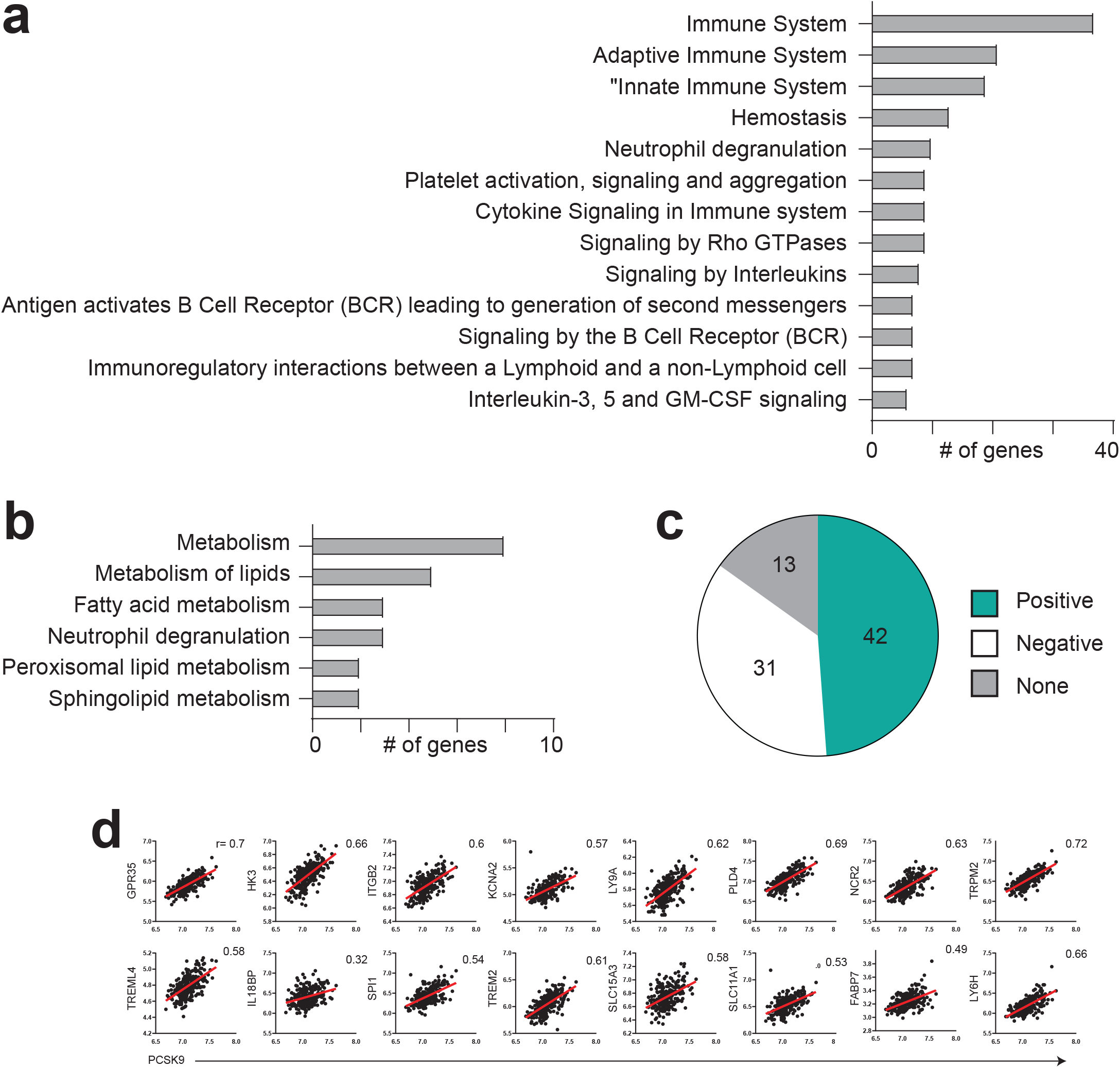
Gene expression patterns unique to each mouse strain and conserved in humans. **a**, Reactome pathways of genes differentially regulated only in the APOE cKO strain at both 10 and 20 weeks after tamoxifen treatment and maintained on a high fat diet throughout. **b**, Reactome pathways of genes differentially regulated only in the D374Y strain at both 10 and 20 weeks after tamoxifen treatment and maintained on a high fat diet throughout. **c**, Venn diagram showing overlap of genes differentially regulated at both 10 and 20 weeks in APOE cKO and D374Y mouse strains that positively correlate with human liver PCSK9 expression. **d**, Examples of selected human liver genes and their correlation with PCSK9 expression. r = Pearson correlation coefficient (Pearsons r). P < 0.05 was considered significant

**Figure S4.**
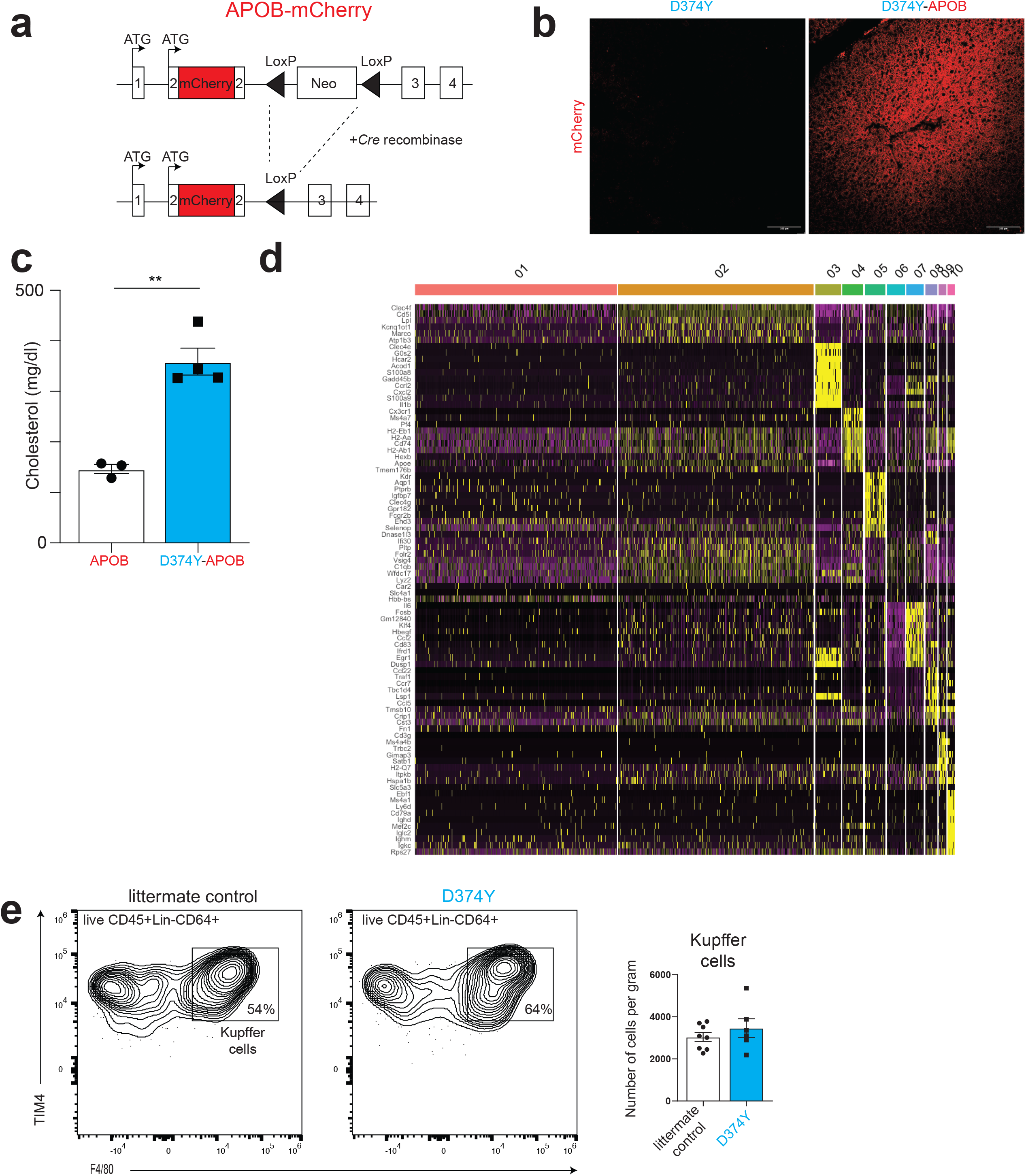
Characterization of mCherry-APOB responsive cells. **a**, Schematic of targeting strategy of the Apob locus to produce mCherry-APOB. **b**, Confocal analysis of paraformalde-hyde-fixed liver from D374Y and D374Y-APOB strains 10 days after tamoxifen treatment and maintained on a chow diet. Scale bars are 100υM **c**, Plasma total cholesterol measurements from female mCherry-APOB and D374Y-mCher-ry-APOB after 4 weeks of high-fat diet following tamoxifen-induction (n = 3-4). Plots are ± SEM, statistical analysis was performed with t-test **p < 0.01. **d**, Heatmap of the top 10 marker genes for every cluster of scRNA-seq data from Live CD45+mCherry-APOB+ cells isolated from liver of female D374Y 10 days after tamoxifen administration and maintained on a chow diet. Yellow indicates high expression of a particular gene, and purple indicates low expression. **e**, Quantification of liver Kupffer cells defined as Live CD45+ Lin-(CD3, CD19, Ly6G) CD64+ F4/80+ TIM4+ in male D374Y and littermate controls 10 days after tamoxifen administration and maintained on a chow diet (n = 5-10).

**Figure S5.**
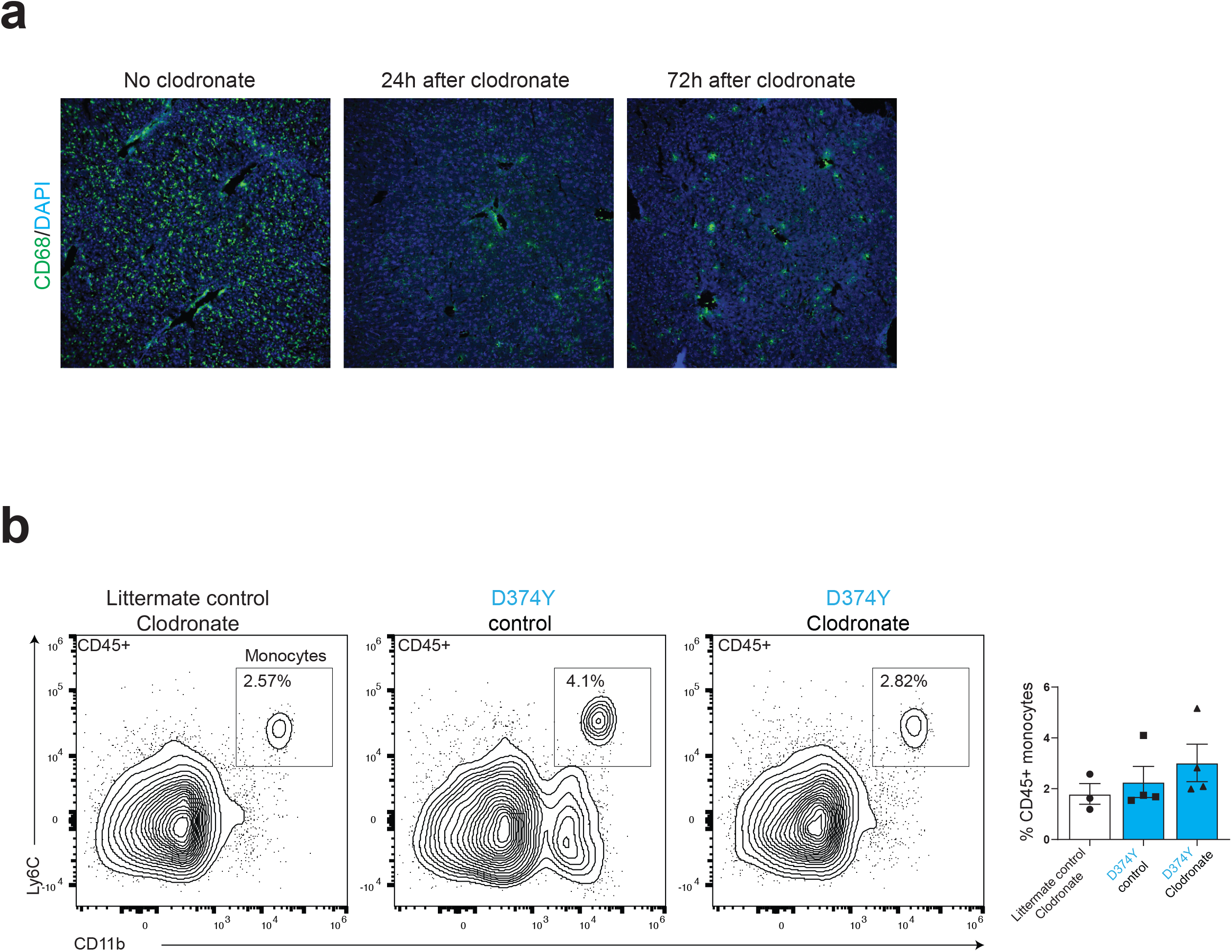
Ablation of Liver KCs but not blood monocytes by clodronate liposomes. **a**, Confocal microscopy of mouse liver before and after clodronate liposome treatment and stained with anti-CD68 and DAPI. **b**, Flow cytometry of blood in female D374Y and littermate controls 10 days after Tamoxifen administration and maintained on a chow diet. Mice were administered clodronate liposome or control Dil liposome as indicated. Blood monocytes were defined as CD45+ CD11b+ Ly6C+ (n = 3-4).

**Figure S6.**
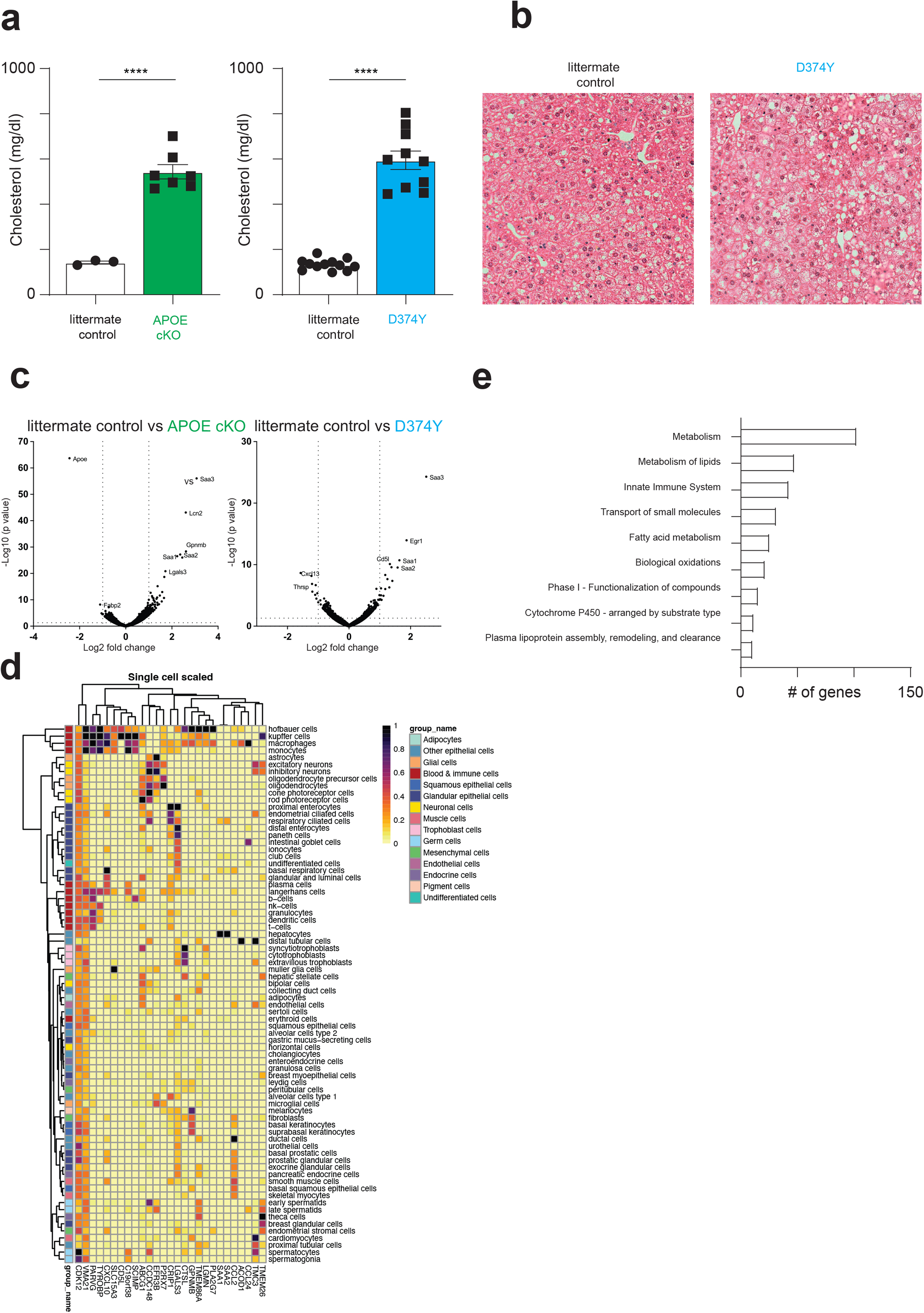
The liver response to four weeks of high-fat diet. **a**, Plasma cholesterol measurements in male and female APOE cKO and female D374Y mice together with their respective littermate controls 4 weeks after Tamoxifen administration and maintained on a HFD (n = 3-12). All plots are ± SEM, Statistical analysis was performed with t-test ****p < 0.0001. **b**, Hematoxylin and eosin staining of paraffin sections of the liver from D374Y and littermate control mice 4 weeks after Tamox-ifen administration and maintained on a HFD. **c**, Volcano plot of differentially expressed genes after 4 weeks of high-fat diet following tamoxifen-induction in APOE cKO or D374Y mice versus respective littermate controls as determined by mRNA-seq. **d**, Expression in 76 human single cell types of the 27 conserved genes from APOE cKO and D374Y alleles 4 weeks after Tamoxifen administration and maintained on a HFD. **e**, Reactome pathways of genes differentially regulated only in littermate control mice after 4 weeks of HFD.

**Figure S7.**
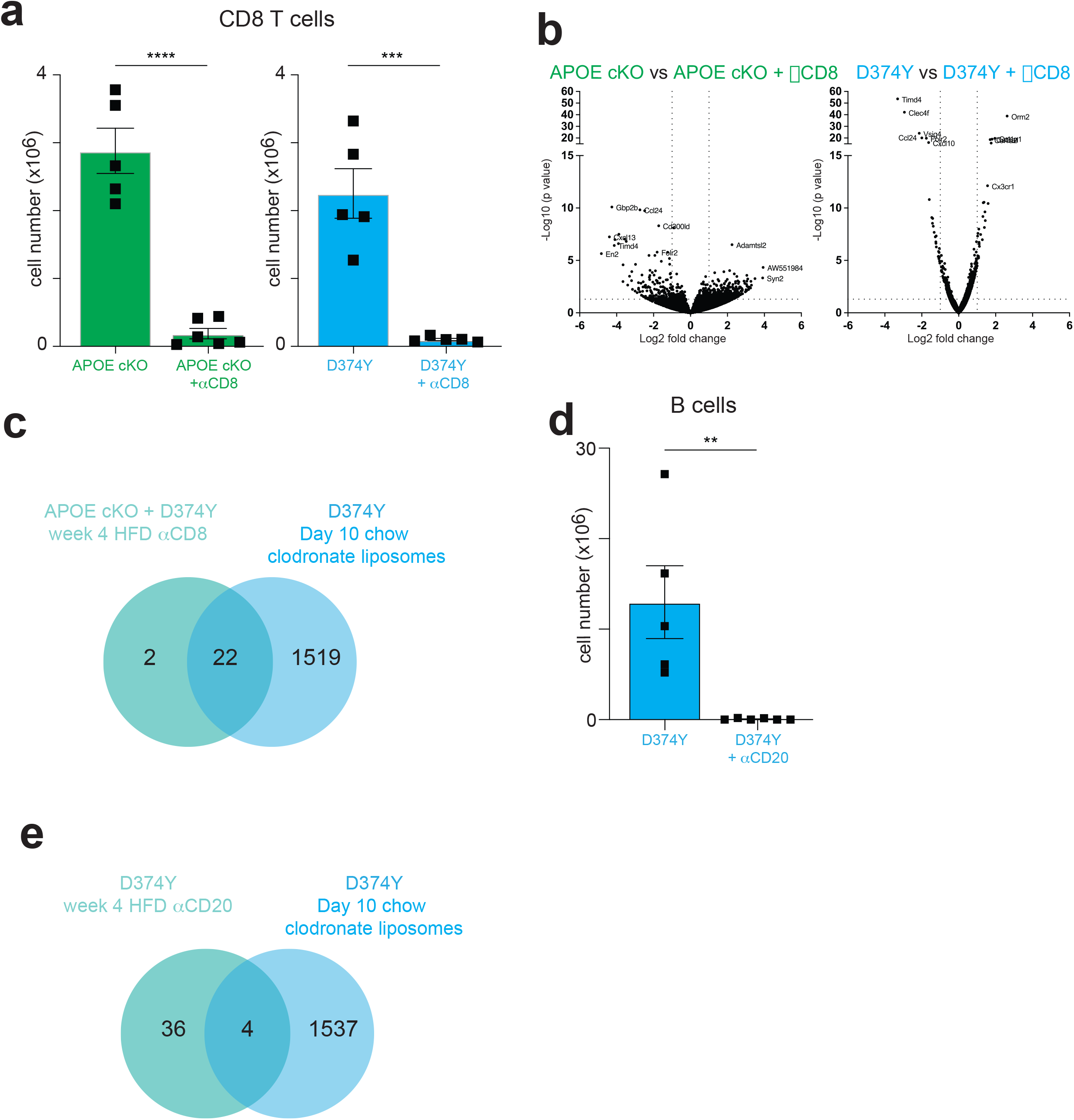
B cells are not required for the KC response to atherogenic dyslipidemia. **a**, Absolute numbers of Spleen CD8 T cells in female in APOE cKO and D374Y mice treated with anti-CD8 antibody relative to PBS treated mice as determined by flow cytometry. Mice were maintained on HFD for 4 weeks following Tamoxifen administration (n = 5-6). **b**, Volcano plot indicating differentially regulated genes in female APOE cKO and D374Y mice treated with anti-CD8 antibody relative to PBS treated mice as determined by mRNA-seq. **c**, Overlap of liver genes differentially regulated in both APOE cKO and D374Y mice depleted of CD8 T cells with clodronate sensitive genes from the day 10 analysis. **d**, Absolute numbers of Spleen B cell (CD19+B220+) in D374Y female mice treated with anti-CD20 antibody relative to PBS treated mice as determined by flow cytometry. Mice were maintained on HFD for 4 weeks following Tamoxifen administration (n = 5-6). **e**, Overlap of liver genes differentially regulated in D374Y ice depleted B cells with clodronate sensitive genes from the day 10 analysis. All plots are ± SEM, Statistical analysis was performed with t-test **p < 0.01, ***p < 0.001, ****p < 0.0001.

